# Immediate Modulation of the Blood Oxygenation Level-Dependent Signals by Dual-Site Transcranial Alternating Current Stimulation Propagates Across the Whole Brain

**DOI:** 10.1101/2024.09.03.610912

**Authors:** Kentaro Hiromitsu, Tomohisa Asai, Hiroshi Kadota, Shu Imaizumi, Masashi Kamata, Hiroshi Imamizu

**Affiliations:** Department of Cognitive Neuroscience, Advanced Telecommunications Research Institute International, Kyoto, Japan; Department of Psychology, The University of Tokyo, Tokyo, Japan; Japan Society for the Promotion of Science, Tokyo, Japan; School of Informatics, Kochi University of Technology, Kochi, Japan; Institute for Education and Human Development, Ochanomizu University, Tokyo, Japan

**Keywords:** Transcranial electrical stimulation, high-definition transcranial alternating current stimulation, functional magnetic resonance imaging, blood oxygenation level-dependent signals, functional brain networks, phase difference

## Abstract

Transcranial alternating current stimulation (tACS) is assumed to target specific brain regions and modulate their activity. Recent discussions of tACS propose that, entraining the phase of brain activity to the stimulation current, stimulation effects extend globally across the whole brain based on phase differences. However, immediate online spatiotemporal propagation of resting-state blood oxygenation level-dependent (BOLD) signals within the brain due to multi-region stimulation remains unclear. The objectives of the present study were three-fold: 1) to elucidate the immediate online effect of tACS on BOLD signal, 2) to examine the extent of the influence on the brain when applying tACS, and 3) to explore whether variations in the phase difference between two brain regions result in differential effects on the stimulated areas and the whole brain. Through two experiments involving high-definition tACS with simultaneous measurements using a functional magnetic resonance imaging (fMRI), we revealed that the immediate online stimulation effects not only altered BOLD signals in the stimulated regions but also propagated across the whole brain in specific spatiotemporal patterns (functional networks). Stimulation effects were observed specifically in regions rich in neural fibres, including the grey and white matter, with no effect in regions containing cerebrospinal fluid. The timing of the signal value peaks depended on the stimulated region and functional networks, with a notable trend observed. Thus, tACS with a specific phase difference in two anatomically connected brain regions can immediately modulate online neural dynamics at both local and global scales.

**Graphical abstract:** 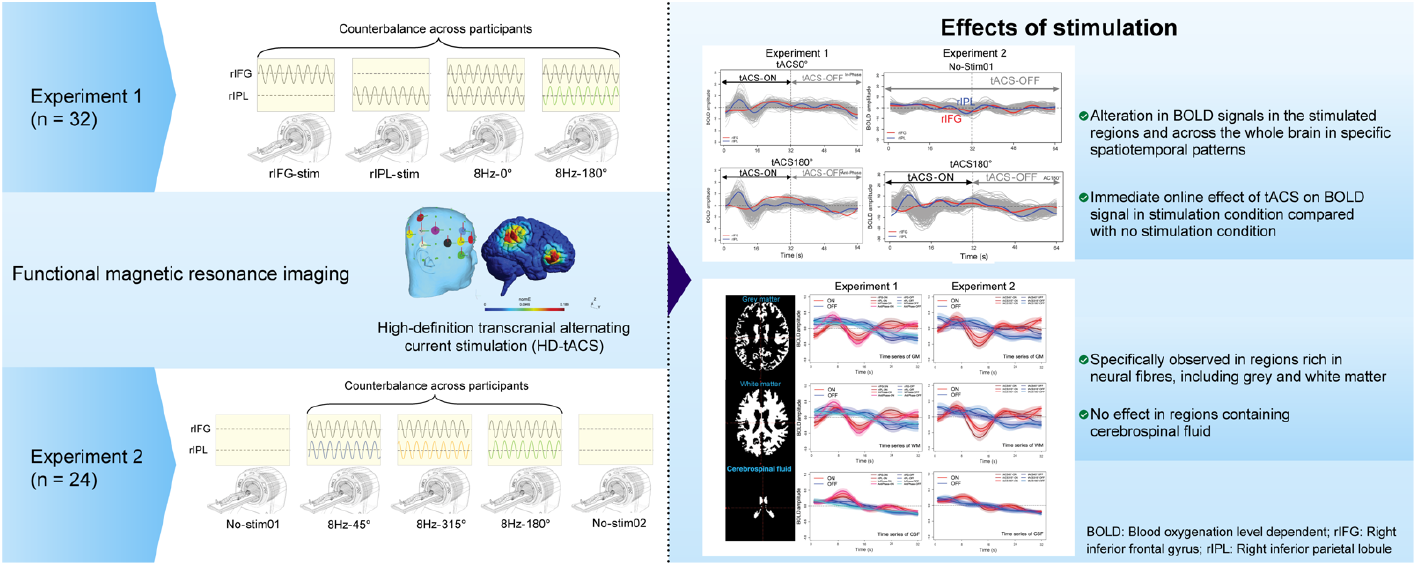

## Introduction

Transcranial alternating current stimulation (tACS) is a one of the non-invasive methods for brain stimulation, and is assumed to target specific brain regions and modulate brain activity. Recent discussions surrounding tACS propose that by leveraging the phase-locking of brain activity to the stimulation current and relying on intra-brain fibres and functional connections, stimulation effects might extend globally across the whole brain with nuances based on phase differences (Wischnewski et al., 2022). tACS is recognised as a non-invasive method for modulating the phase of neural activity in specific brain regions, wherein an alternating current is applied between two or more electrodes placed on the scalp to modulate the phase difference between the regions. Neurophysiological findings in primates suggest that tACS can alter the neural spike timing, inducing local online neural entrainment (Johnson et al., 2020; Krause et al., 2019, 2022). Furthermore, alterations in neurophysiological mechanisms, such as the synaptic plasticity or long-range connectivity, which are not limited to neural spike timing alterations alone, have been discussed (Wischnewski et al., 2022). In terms of its effects on cognitive function, distinct intervention effects of tACS based on the previously described neurophysiological mechanisms have been reported (Grover et al., 2022; Reinhart & Nguyen, 2019). The effects of tACS on neural activity and cognitive function have been demonstrated in this manner, but there remain three unresolved issues when examining the relationship between brain activity and cognitive function using functional magnetic resonance imaging (fMRI). 1) The duration of the effects induced by tACS (temporal domain). 2) The extent to which the effects of tACS spread across the brain and consequently lead to changes in neural activity (spatial domain), and 3) The lack of clarity regarding the optimal phase difference when applying tACS to multiple brain regions (phase domain).

First issue is relating to whether tACS exerts its effects only when delivered online, offline (i.e. after-effects), or both. Several studies have shown that online effect of tACS on behaviours (Grover et al., 2022; Helfrich et al., 2014), and on electrophysiology (e.g., TMS-induced MEP, Antal et al., 2008; Pozdniakov et al., 2021). However, it has also been reported that the size of motor evoked potentials (MEP) induced by transcranial magnetic stimulation (TMS) remains consistently enhanced even after tACS stimulation, indicating an offline effect (Guerra et al., 2020). Furthermore, the confirmation of both online and offline effects has been reported depending on the task and the parameters of tACS (Fujiyama et al., 2023). It has been hypothesised that local online entrainment causes neural plasticity, as a result, online effect leads to the after-effect (Vossen et al., 2015).

However, the debate surrounding these online and offline effects has primarily been discussed based on task-related metrics, and particularly in neurophysiological studies of the human brain using fMRI, the effects of tACS on resting-state blood oxygenation level-dependent (BOLD) signals remain unclear. For instance, while Antal et al. (2011) demonstrated that task-related BOLD signals are modulated by tDCS, they reported no changes in resting-state BOLD signals. Addressing such findings, Vosskuhl et al. (2016) suggested that changes in resting-state fMRI BOLD signals may be too small to detect compared to task-related signals or may be undetectable by fMRI. However, animal studies have reported immediate changes in blood flow under anaesthesia with tACS (Turner et al., 2021) and burstiness (changes in neuronal spike frequency within a certain period) during awake states in monkeys (Johnson et al., 2020), suggesting the potential for immediate online effects to be measured by fMRI even during human resting states. Therefore, in this study, to investigate immediate online effects of tACS in human resting-state fMRI, short trains of tACS were repeated, paired with baseline trains without stimulation (Vosskuhl et al., 2016).

Secondly, regarding the structural impact of stimulation on specific regions of the brain, the prevailing notion posits that tACS elicits localised effects, thereby influencing the modulation of brain function. However, recent findings suggest that the impact of tACS extends beyond the stimulated brain region, thereby influencing broader brain networks (Wischnewski et al., 2022). In human tACS research, concurrent measurements with fMRI or electroencephalography (EEG) have confirmed that brain activity in several regions or networks according to the stimulated regions are affected (Violante et al., 2017; Vossen et al., 2015; Vosskuhl et al., 2016). This issue arises from the use of large electrodes in electrical stimulation (e.g., 5 × 7 cm) and the necessity to place not only the target electrode, but also the reference electrode, in a distant region when stimulating specific areas. To address this issue, high definition tACS (HD-tACS) has been developed, enabling spatially precise stimulation of the target area. HD-tACS is characterised by surrounding stimulation electrodes with oppositely polarised return electrodes, and employs smaller electrodes than conventional methods (Wu et al., 2021). However, even with HD-tACS, variations in the montage of the stimulation electrodes and stimulation parameters lead to variability in the observed effects in the brain. This is attributed to the possibility that the effects of tACS on specific neurons spread across a broad range of the brain (i.e. travelling waves; Muller et al., 2018), with varying propagation patterns depending on the stimulated region. However, the extent to which tACS, including HD-tACS, affect the brain, remains unclear. To elucidate this, the current study not only focuses on changes in BOLD signals in regions stimulated by tACS, but also examines anatomical subdivisions (such as grey matter [GM], white matter [WM], and cerebrospinal fluid [CSF]) to investigate the extent of tACS effects within the brain. Considering the debated possibility of tACS effects extending to deep brain regions such as the hippocampus and insula (Shan et al., 2023), along with the phenomenon of neural entrainment to tACS frequencies (Johnson et al., 2020), it is predicted that the effects of tACS may extend widely to regions rich in neurons and neural fibres, including both grey and white matters.

Lastly, in the application of multi-electrode stimulation, researchers determine the optimal phase differences that influence brain activity and cognitive function. Previous studies have commonly employed binary-phase manipulations involving in-phase and anti-phase stimulations (Polanía et al., 2012). In-phase stimulation, characterised by a stimulation current phase difference of 0° between two brain regions, is anticipated to enhance behavioural performance by synchronising activity in the respective areas of the brain. Conversely, anti-phase stimulation, that is, a phase difference of 180°, is expected to exert an opposite effect to in-phase stimulation. For example, Reinhart & Nguyen (2019) revealed that frontotemporal theta phase synchronisation (in-phase stimulation), but not desynchronisation (anti-phase stimulation), leads to improved working memory in older adults. However, simulation studies have disputed the biophysical validity of the binary effects of phase differences (Saturnino et al., 2017).

Intracranial neural recordings of multi-electrode tACS effects on the primate brain have demonstrated an in-vivo nonlinear relationship between the phase difference of the applied currents and both the phases and magnitudes of intracranial electric fields (Alekseichuk et al., 2019). These results imply that the influence of phase differences may vary depending on the neurophysiological features of the specific brain regions being stimulated. Given the inherent regional lags in brain activity, as exemplified by the phenomena of travelling waves (Bolt et al., 2022), it is plausible to posit that the phase differences amenable to intervention via tACS may vary across brain regions.

In light of the unresolved issues surrounding the immediate online effect of tACS during resting-state fMRI, the specific reach of tACS impact on distinct brain regions, and the modulation of effects through inter-regional stimulation phase differences, the current study aimed to achieve three primary objectives: 1) to elucidate the immediate online effect of tACS by comparing the short-term BOLD signals during tACS with periods where no stimulation is applied, 2) to examine the extent of the influence on the brain when applying tACS by comparing the effects on BOLD signals across anatomical subdivisions, and 3) to explore whether variations in the phase difference between the two stimulated regions result in differential effects on the BOLD signals in stimulated areas and across the whole brain. We targeted the frontal and parietal areas of the right hemisphere of the brain because these two regions are connected anatomically (SLF; Thiebaut de Schotten et al., 2011) and functionally (Mars et al., 2012) and are considered critical for several cognitive functions (e.g., working memory) (Violante et al., 2017). We anticipate that the optimal phase difference causes a greater change of the BOLD signals in the region connecting the two areas, compared with that in the non-optimal or control conditions. In addition, because the inherent lag in brain activity is a natural phenomenon, we expect to observe a spread based on the brain structures or networks.

## Methods

### Participants

We enrolled a total of 56 participants, of whom 32 were exposed to Experiment 1 (mean age 22.06 ± 1.78, with a standard deviation of 1.78, including 12 females) and 24 to Experiment 2 (mean age 22.13 ± 2.42, with a standard deviation of 2.42, including 5 females); all of whom were right-handed. Prior to participation, written informed consent was obtained from each participant, following the procedures approved by the Office for Life Science Research Ethics and Safety at The University of Tokyo (Ethics Review Number: 20-42). This study was conducted in accordance with the principles of the Declaration of Helsinki.

To ensure the safety of the participants, a comprehensive checklist for fMRI and tACS safety was accomplished by all participants before engaging in the experiment. All participants confirmed the absence of a history of neurological disorders, mental illness, head injuries, or metal implants or devices in either the body or head; reported no current use of psychoactive medication; and confirmed non-pregnancy.

### HD-tACS

Alternating current stimulation was administered noninvasively using a magnetic resonance-compatible multichannel stimulator (DC-Stimulator MC; NeuroConn GmbH, Ilmenau, Germany). The stimulation waveform was sinusoidal, with a frequency of 8 Hz, a peak-to-peak amplitude of 4 mA, and 1 s ramp-up and ramp-down intervals. Circular rubber electrodes with a diameter of 2 cm were used. Impedance recorded within the fMRI room was maintained below 15k ohm, facilitated by the application of a conductive paste (Ten20, Weaver & Company, Aurora, CO, USA).

The target regions for stimulation were the right inferior frontal gyrus (rIFG) and right inferior parietal lobule (rIPL). Electrodes were strategically placed according to the International 10-20 System: one electrode was positioned at the centre of the target region (F8 for rIFG and CP6 for rIPL), whereas the remaining four electrodes surrounded the target region (F6, FT8, AF8, and F10 for rIFG; P6, C6, CP4, and TP8 for rIPL) (Figure 1). The configuration of these electrodes for each target region was determined using electric field simulations with SimNIBS 3.2 (Saturnino et al., 2018). Under stimulation conditions, stimulations occurred for 32 s (tACS-ON), followed by a 32-s period without stimulation (tACS-OFF); this cycle was repeated five times, totalling 320 s for each condition (Figure 2).

**Figure 1.**
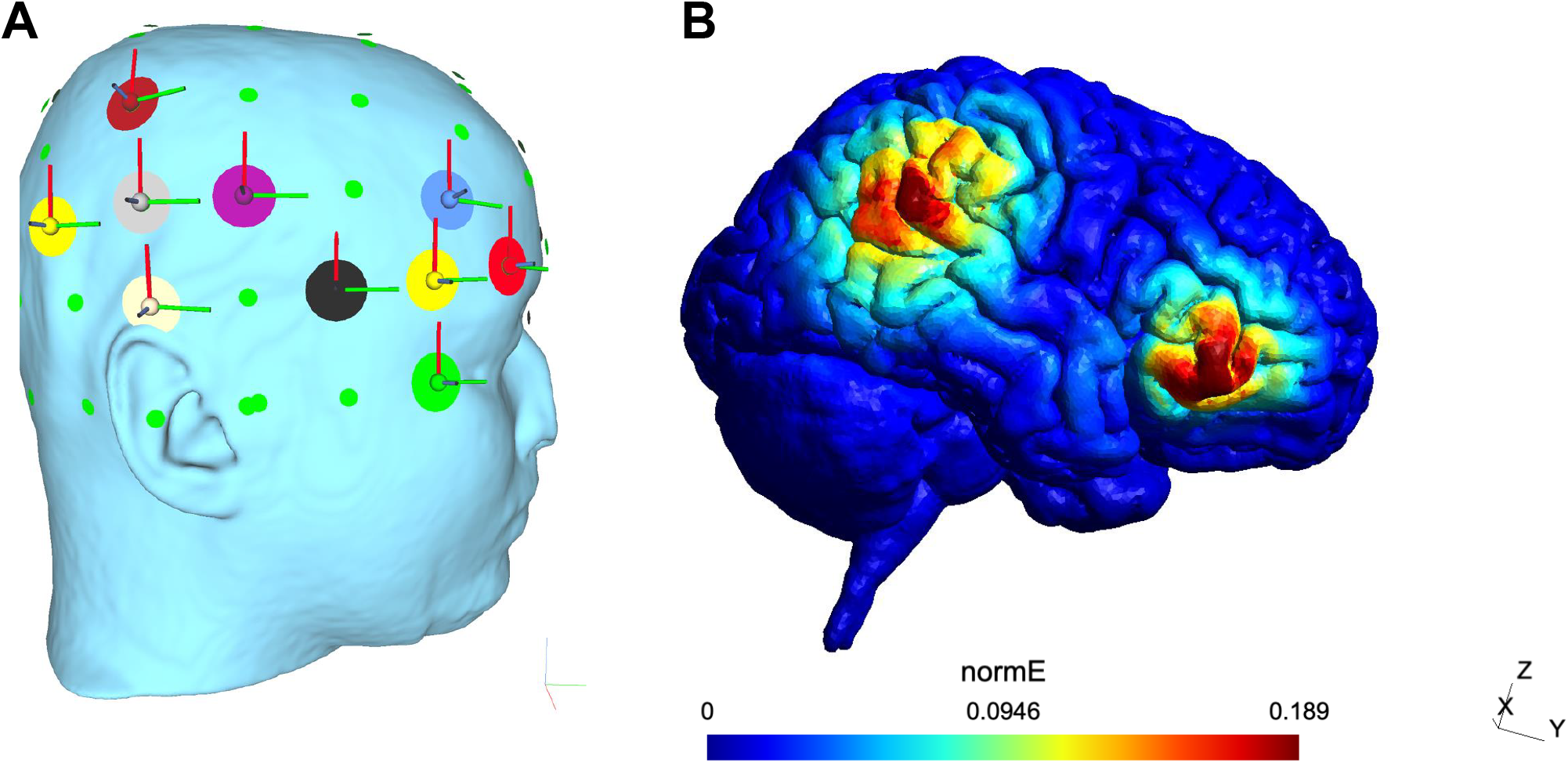
Electrode montage and the simulation of electric field generated by frontoparietal tACS. (A) The target regions for stimulation were the rIFG and rIPL. This figure illustrates concurrent stimulation of the two regions on the right side of the brain. As a montage, one electrode was positioned at the centre of the target region (F8 for rIFG; CP6 for rIPL), while the remaining four surrounded the target region (F6, FT8, AF8, and F10 for rIFG; P6, C6, CP4, and TP8 for rIPL). (B) Electric field simulation was done using SimNIBS 3.2 (Saturnino et al., 2018). tACS, transcranial alternating current stimulation; rIFG, right inferior frontal gyrus; rIPL, right inferior parietal lobule.

**Figure 2.**
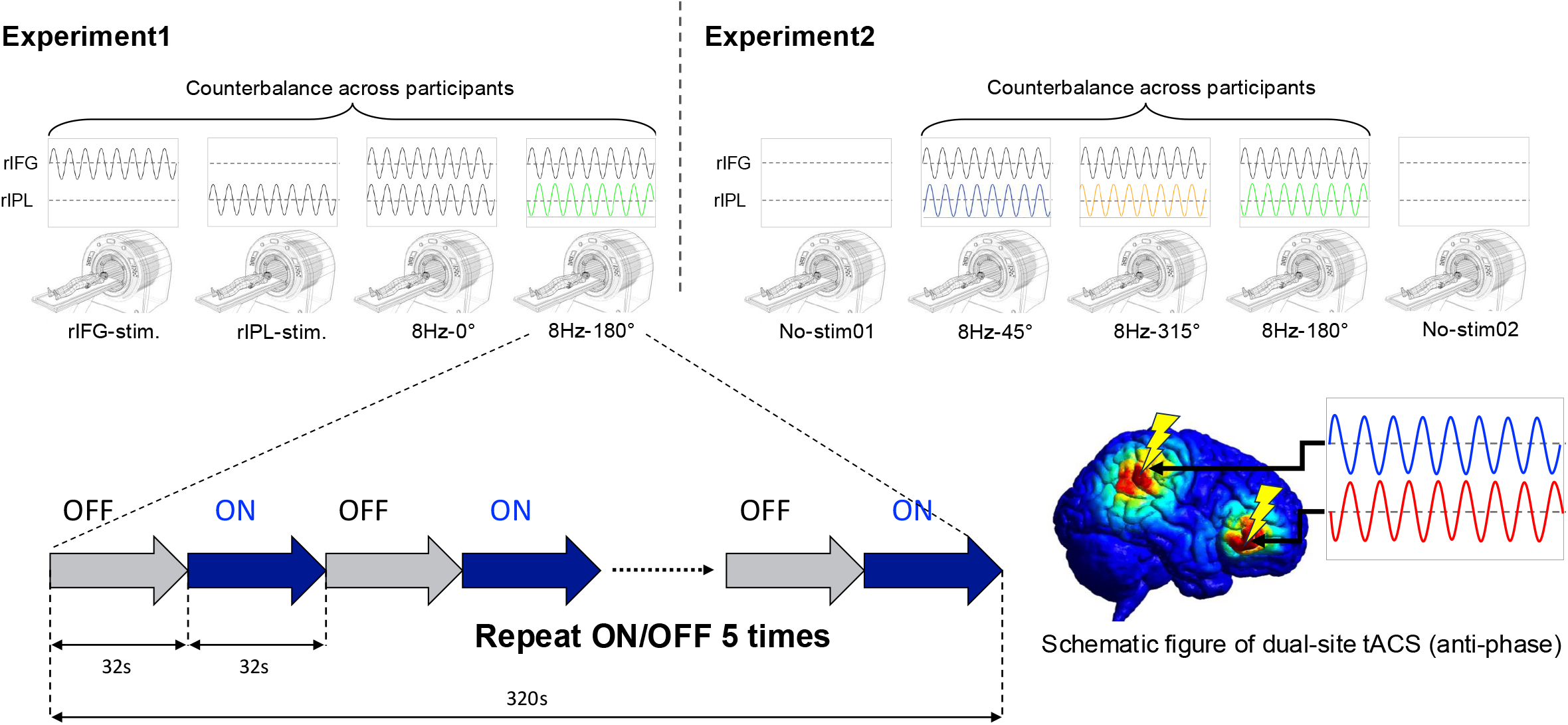
Procedure for Experiments 1 and 2. In Experiments 1 and 2, fMRI was utilised during the application of 8 Hz tACS targeting the rIFG and rIPL. Five alternating epochs of tACS-ON (32 s) and tACS-OFF (32 s) were done, with counterbalancing starting from tACS-ON or -OFF across subjects. In Experiment 1, four tACS conditions were employed, whereas Experiment 2 employed three tACS conditions with resting-state imaging performed before and after tACS. The order of the tACS conditions was counterbalanced across subjects in both Experiments 1 and 2. The schematic at the bottom right illustrates the condition for stimulating the rIFG and rIPL with a phase difference of 180°. fMRI, functional magnetic resonance imaging; tACS, transcranial alternating current stimulation; rIFG, right inferior frontal gyrus; rIPL, right inferior parietal lobule.

In Experiment 1, tACS was applied across four distinct conditions: 1) rIFG stimulation; 2) rIPL stimulation; 3) in-phase stimulation, where the phase difference between electrical currents for rIFG and rIPL was 0°; and 4) anti-phase stimulation, where the phase difference between electrical currents for rIFG and rIPL was 180°. Stimulation under conditions 1 and 2 exclusively targeted a single region (i.e., the rIFG or rIPL), while stimulation under conditions 3 and 4 targeted both regions with a specific phase difference (0° or 180°).

In Experiment 2, tACS was applied across five different conditions: 1) initial no-stimulation condition conducted prior to the rest of the stimulation conditions (No-Stim01); 2) final no-stimulation condition conducted after completing all stimulation conditions (No-Stim02); 3) 45° phase difference condition; 4) 180° phase difference condition; and 5) 315° phase difference condition. Stimulation in conditions 3–5 simultaneously targeted both the rIFG and rIPL.

### Procedures

Each experiment consisted of four or six tACS conditions. Throughout the resting-state fMRI scans, sequences of tACS-ON and tACS-OFF epochs were iterated five times, each lasting 32 s. The order in which tACS-ON preceded tACS-OFF, or vice versa, was counterbalanced across participants. The duration of each run was 320 s. In Experiment 1, four tACS conditions, each comprising five repetitions of 32-s tACS-ON and 32-s tACS-OFF, were administered in a randomised order. In Experiment 2, three tACS conditions, excluding the initial and final sequences (i.e., sequences without tACS), were administered in a randomised order (Figure 2). During the resting-state fMRI scans, participants were provided with the instruction to “Relax. Stay Awake. Fixate on the central cross mark and refrain from concentrating on specific things.”

### MRI acquisition

Resting-state fMRI and T1-weighted (T1w) structural MRI data were systematically acquired from all participants using a 3T Siemens MAGNETOM Prisma scanner at the IRCN Human fMRI Core, University of Tokyo, equipped with a 32-channel head coil. Resting-state functional images were obtained using a gradient-echo multi-band echo-planar imaging sequence with the following parameters: repetition time (TR), 0.8 s; echo time, 34.4 ms; flip angle, 52°; field of view, 206 mm; matrix dimensions, 86 × 86; and voxel resolution, 2.4 mm. Each run comprised 420 volumes, and the initial 20 scans were discarded to ensure signal stability. For susceptibility distortion correction of the resting-state functional images, a main magnetic field map was acquired with the following specifications: TR, 6.1 s; echo time, 60 ms; flip angle, 90°; field of view, 100 mm; matrix dimensions, 64 × 64; and voxel resolution, 2.4 mm.

### fMRI data preprocessing

We performed a preprocessing pipeline for resting-state fMRI using fMRIPrep version 20.2.0 (Esteban et al., 2019). The first 20 scans were discarded to ensure signal stability. The preprocessing steps included realignment, co-registration, distortion correction with a field map, segmentation of the T1w structural images, motion correction, and normalisation to the Montreal Neurological Institute (MNI) space.

To enhance the specificity of our analysis for low-frequency fluctuations indicative of resting-state fMRI BOLD signals, a temporal bandpass filter was implemented on the time series. This filter, employing a Butterworth filter with a pass band ranging between 0.01 Hz and 0.1 Hz, served to refine the data. Additionally, we applied spatial smoothing to the temporally filtered fMRI data by employing a Gaussian kernel with a full width at half maximum of 5 mm. These preprocessing steps collectively ensured the robustness and appropriateness of the resting-state fMRI data for subsequent analyses.

### fMRI data analysis

After preprocessing, the BOLD signal time series was extracted from 264 regions of interest (ROIs) (Power et al., 2011). The ROIs of the stimulated regions were 5 mm-width spheres created based on the MNI coordinates of the regions directly beneath the stimulation electrode. In each stimulation condition for each participant, each ROI’s BOLD signal time series of the stimulation (tACS-ON) and no-stimulation (tACS-OFF) epochs were summed and averaged respectively in order to compare the effect of tACS on the BOLD signal. Furthermore, to compare the effects at a global level within the brain, the whole brain was divided into three regions: GM, WM, and CSF, and the average BOLD signal time series was extracted from each region. To investigate the extent of the signal changes induced by tACS from baseline in the BOLD signal time series for each ROI, we performed baseline correction after re-referencing. Initially, the time points of the time series were up-sampled by a factor of ten using spline interpolation. The mean of each ROI’s entire time series for each stimulation condition was subtracted from the corresponding time series (referencing). Further processing involved subtracting the mean of 50, 100, and 200 scans in the preceding tACS-OFF epoch from each tACS-ON-OFF epoch (baseline correction) (Tanner et al., 2016).

## Results

### Immediate online modulation of BOLD signals in the stimulated regions through tACS

To examine the effects of tACS on both stimulated regions, we computed the cumulative averages of the tACS-ON, tACS-OFF, and tACS-ON-OFF epochs for each stimulation condition in Experiments 1 and 2. In the tACS-ON epochs, an immediate change in BOLD signals was evident within the stimulated region (Figure 3). Regarding the tACS-OFF epochs, there was an absence of signal elevation, with a discernible typical and relatively gradual fluctuation apparent during resting-state fMRI. To examine the immediate online effect of tACS, the BOLD signal between tACS-ON and -OFF epochs across time for each stimulation condition was compared (Figure 3A and C). In Experiment 1, two-way repeated measures analysis of variance (ANOVA) revealed that in the rIFG region under both single-site and two-site stimulation conditions, the BOLD signal significantly increased during the latter half of the tACS-ON epoch compared to the tACS-OFF epoch. For the rIPL region, during the first half of the tACS-ON epoch only in the tACS 180° condition, there was a significantly higher BOLD signal compared to the tACS-OFF epoch. In Experiment 2, in the rIFG region, only under the tACS 45° condition during the first half of the epoch, the signal values of the tACS-ON epoch were significantly lower than those of the tACS-OFF epoch. Conversely, under the tACS 135° and tACS 180° conditions, the tACS-ON epoch exhibited higher signal values compared to the tACS-OFF epoch during mid-epoch. In the rIPL region, the waveforms across conditions were similar, however, only under the tACS 180° condition, the tACS-ON epoch exhibited significantly higher signal values compared to the tACS-OFF epoch during the latter half of the epoch. On the other hand, the substantial increase in BOLD signal values was not observed in the two tACS-OFF resting-state conditions (Figure 3C). There were no significant differences between signals in Pseudo-ON and Pseudo-OFF epochs under both No-Stim01 and No-Stim02 conditions.

**Figure 3.**
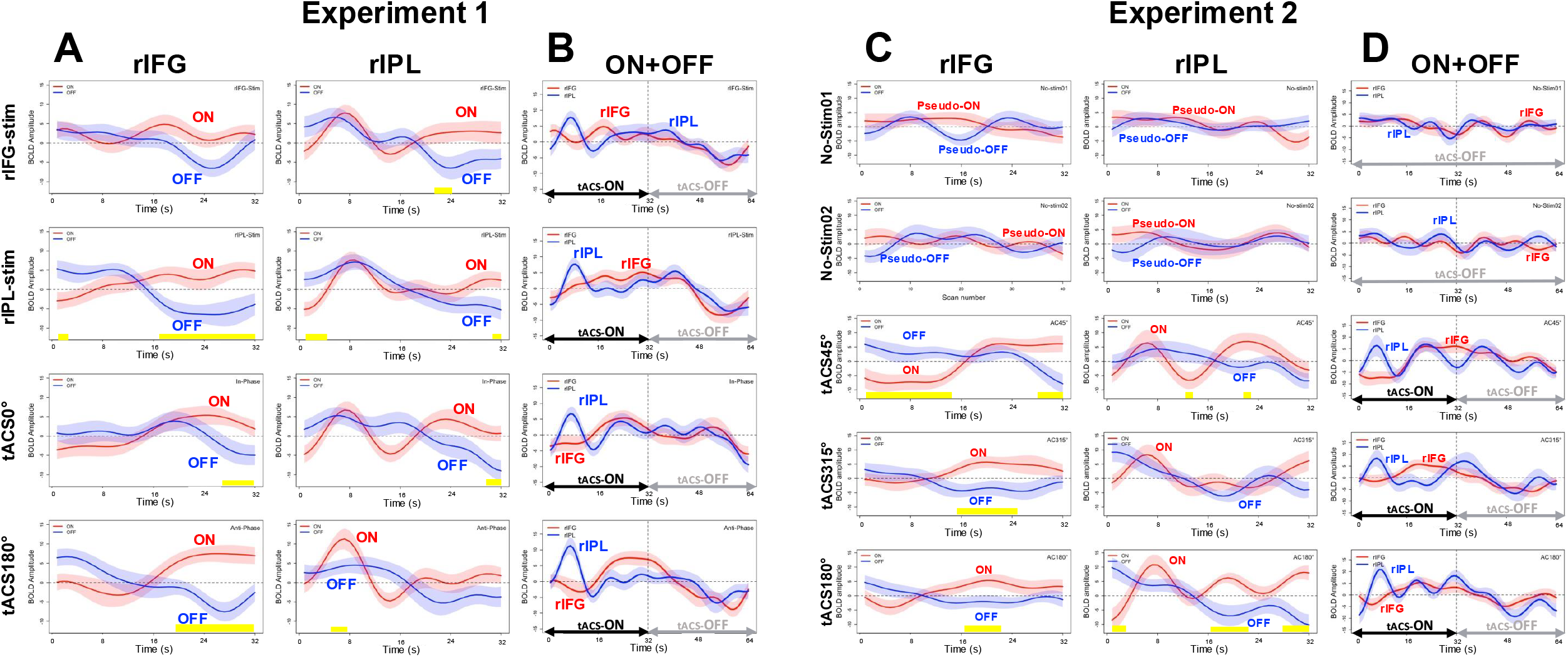
Modulation of BOLD Signals within tACS-Targeted Regions. The left side of the figure illustrates the tACS-induced changes in BOLD signals of the stimulated regions during Experiment 1, whereas the right side shows those in Experiment (A) and (C) present the epoch-wise averaged results for the tACS-ON (red line) and -OFF (blue line) conditions in each experiment in the stimulated regions (rIFG and rIPL). The yellow rectangles indicate the time course of significant interaction between the presence of tACS and time in the ANOVA, as well as significant simple main effects of the presence of tACS per time. The BOLD signals, pseudo-aligned based on the tACS-ON and -OFF orders of the tACS conditions for each participant, are depicted together with their averaged BOLD signals (Pseudo-ON and Pseudo-OFF). (B) and (D) show the average BOLD signals across consecutive tACS-ON and -OFF epochs, with the red and blue lines denoting the rIFG and rIPL regions, respectively. The ribbons in each graph delineate mean ± SE. BOLD, blood oxygenation level-dependent; tACS, transcranial alternating current stimulation; rIFG, right inferior frontal gyrus; rIPL, right inferior parietal lobule; fMRI, functional magnetic resonance imaging; SE, standard error.

Regarding the behaviour of the BOLD signal, the rIPL ROI time series suggested a surge, characterised by prompt elevation, proved to be transitory, and displayed a characteristic trend known as a haemodynamic response function. When comparing tACS conditions, it was evident that the tACS 180° condition exhibited the most substantial increase in signal values. Conversely, in the time series of the rIFG ROI, a gradual decline in signal values was first observed, with the exception of the rIPL stimulation condition in which no stimulation was applied to the rIFG. Following this initial decline, a gradual increase in signal values was observed after 16 s. Thus, contrasting features were observed, wherein an immediate surge in signal values in rIPL and a gradual increase of signal values in rIFG following an initial decline was observed. Temporal changes in BOLD signals across the whole brain, segmented by a Power ROI, are shown in the Supplementary Animation. Upon visual inspection, a notable increase in signal values was observed surrounding the stimulated regions of the rIFG and rIPL. During the tACS-ON epochs, a more pronounced elevation in signal values was observed in the region between these two stimulated areas than in other regions. Nevertheless, alterations in brain activity were evident not only within the targeted regions following tACS but also across the whole brain. In contrast, during unstimulated resting-state fMRI, a sequential and seemingly random pattern of changes was observed.

### Baseline correction for BOLD signals time series

The increase in BOLD signals induced by tACS is immediate, but when compared to the results of the resting state without stimulation (Figures 3), it was observed that specific oscillations persist for a certain period, even after the effect on signal decay. In other words, based on the methodology of this study involving the repetition of tACS-ON and tACS-OFF epochs, the onset of the tACS-ON epoch, unlike the actual resting state, may still exhibit the residual effects of tACS, albeit to some extent. Therefore, we applied a baseline correction by subtracting the average of several scans in the tACS-OFF epoch from that in the tACS-ON epoch to align the onset of stimulation with the baseline.

As depicted in Figure 4, a baseline correction was applied, aligning the starting point of the tACS-ON epoch to zero for all conditions. The baseline was defined by subtracting the mean of 10 scans in the preceding tACS-OFF epoch from each tACS-ON-OFF epoch. The length of the tACS-OFF epoch used as the baseline was determined by considering the frequency of the BOLD signal. To examine the change from the baseline in each stimulated region, corrected BOLD signal values were compared with zero using one-sample *t*-test with multiple comparison correction using false discovery rate (FDR) (Figure 4). The result suggested that, in Experiment 1, the BOLD signal in the rIFG ROI under the tACS 180° condition showed a significant increase from the baseline during the latter half of the tACS-ON epoch (*ps* < 0.05, FDR). However, the BOLD signal from the baseline in the rIFG and rIPL ROIs under other tACS-ON conditions showed no significant changes. In Experiment 2, it was found that the BOLD signals in the rIFG ROI under the tACS 45° and 180° conditions exhibited significant changes from the baseline during the initial signal decrease of the tACS-ON epoch (*ps* < 0.05, FDR). Additionally, the BOLD signals in the rIPL ROI showed a significant increase from the baseline during the first half of the tACS-ON epoch under the tACS 180° condition (*ps* < 0.05, FDR). The BOLD signals of other tACS conditions in both rIFG and rIPL ROIs did not show a change from the baseline. In the absence of stimulation during the resting-state condition (i.e., No-Stim01 and No-Stim02), the BOLD signal fluctuated around zero.

**Figure 4.**
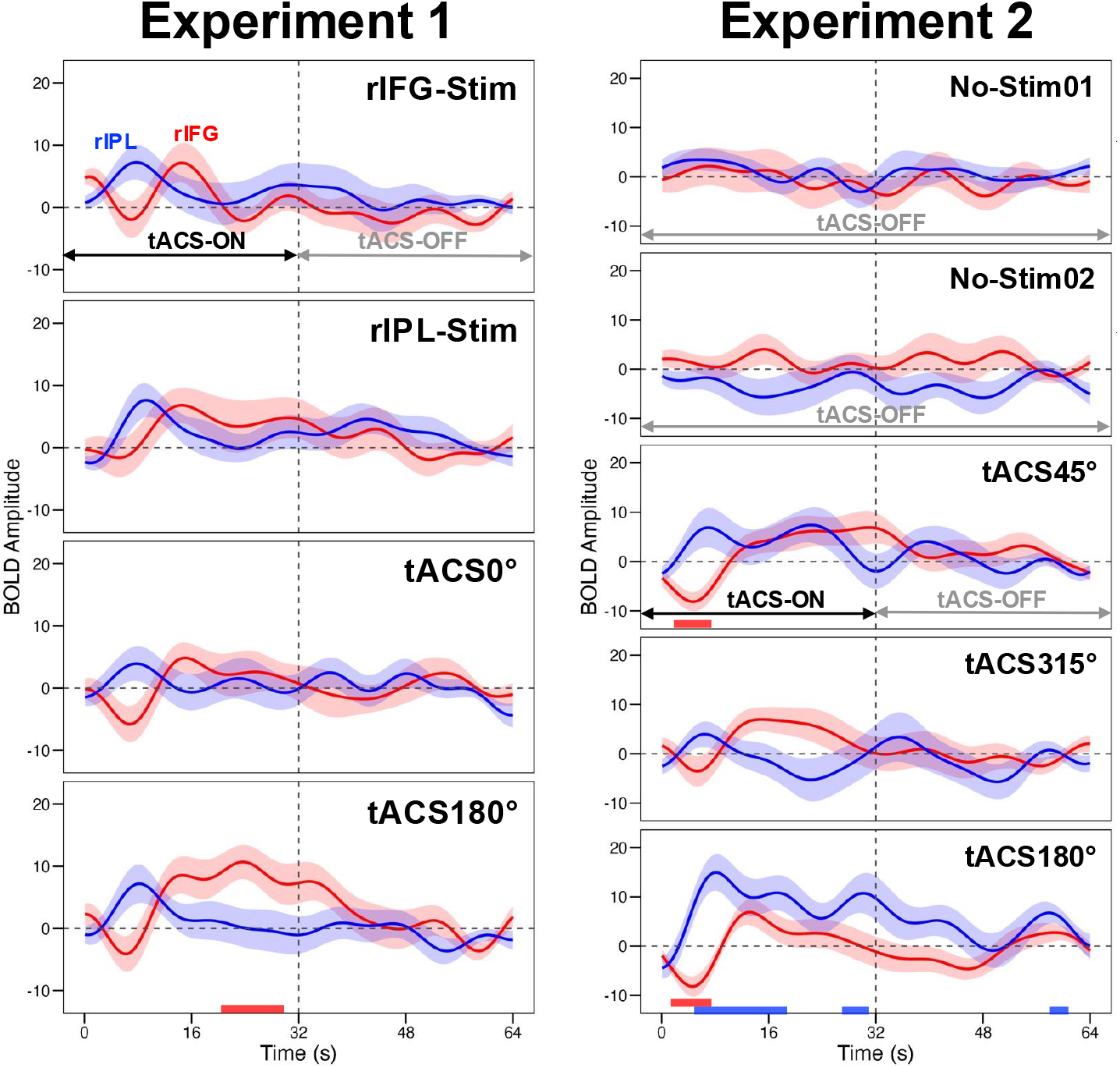
BOLD Signals after Baseline Correction. The left panel of the figure depicts the BOLD signals originating from the stimulated regions in Experiment 1. The right panel shows the corresponding signals from Experiment 2. Each time series represents the average BOLD signal after baseline correction across successive epochs of tACS-ON and -OFF across subjects. The red and blue rectangles indicate the periods of significant difference from zero (*ps* < .05, FDR) for each stimulated region. BOLD, blood oxygenation level-dependent; ROI, region of interest; tACS, transcranial alternating current stimulation.

### Effects of tACS on global brain regions: GM, WM, and CSF

As shown in Figure 5, an increase in BOLD signals induced by tACS was observed not only in the stimulated regions but also across a broad area of the brain. As shown in Figure 5, the BOLD signals originating from the stimulated regions and the whole brain as defined by the Power ROI (comprised of 264 nodes) showed an immediate change in BOLD signals not only within the stimulated regions but also across the whole brain under all stimulation conditions. Subsequently, we investigated the extent to which the effects of tACS extend beyond the stimulated regions (Figure 6A). To elucidate the extent of such effects throughout the brain, we examined temporal changes in BOLD signals (subtraction between minimum value from maximum value) for each global anatomical subdivision of the brain (Figure 6B).

**Figure 5.**
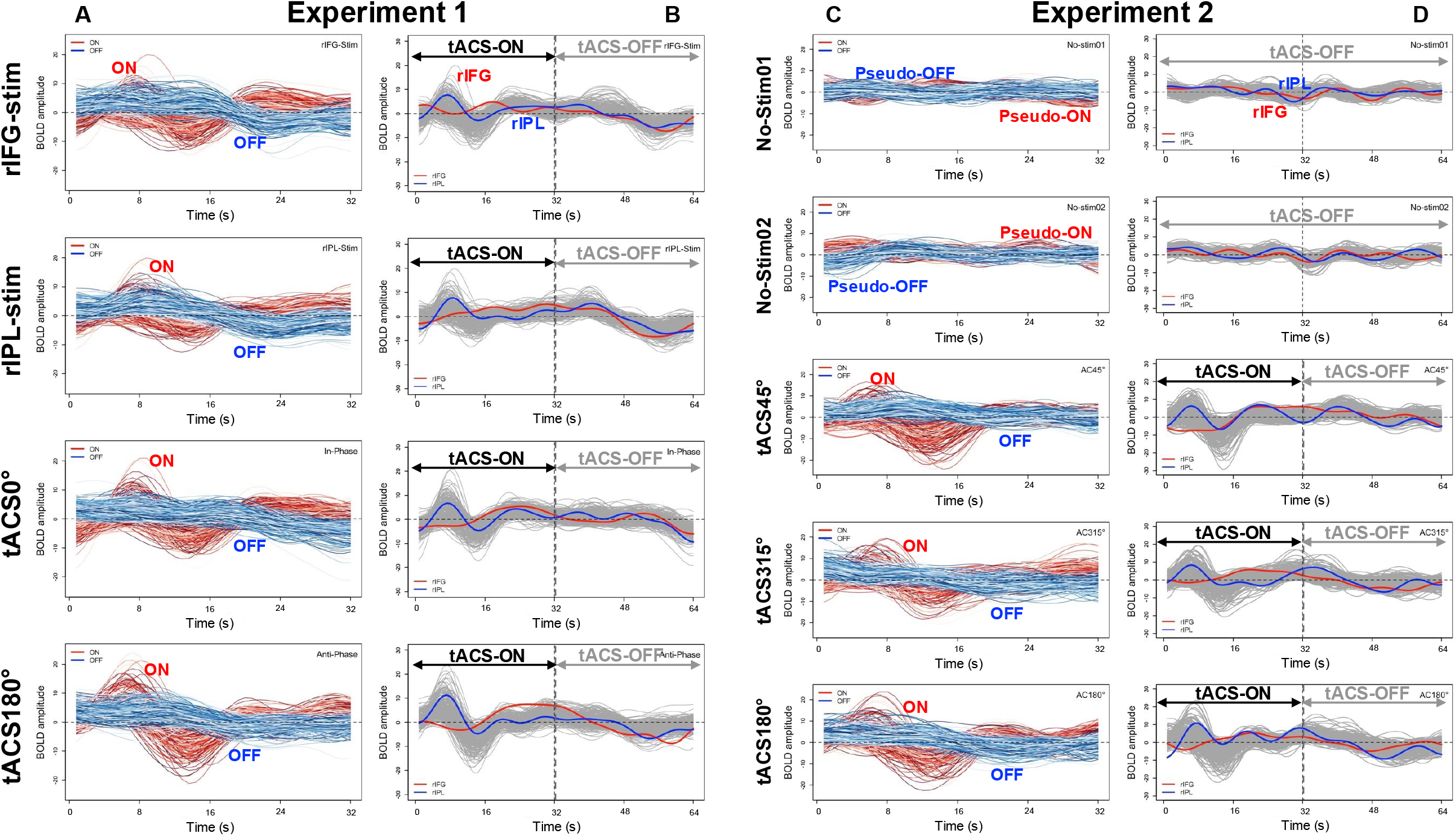
Modulation of BOLD Signals across the Whole Brain. (A) and (C) depict the results of averaging the BOLD time series of each ROI for each tACS-ON (red line) and -OFF (blue line) epoch using Power ROI (Power et al., 2011), which consisted of 264 ROIs. (B) and (D) illustrate the averaged BOLD signals across consecutive tACS-ON and -OFF epochs of the whole brain (grey lines); the red line denotes the rIFG region, and the blue line represents the rIPL region. BOLD, blood oxygenation level-dependent; ROI, region of interest; tACS, transcranial alternating current stimulation; rIFG, right inferior frontal gyrus; rIPL, right inferior parietal lobule.

**Figure 6.**
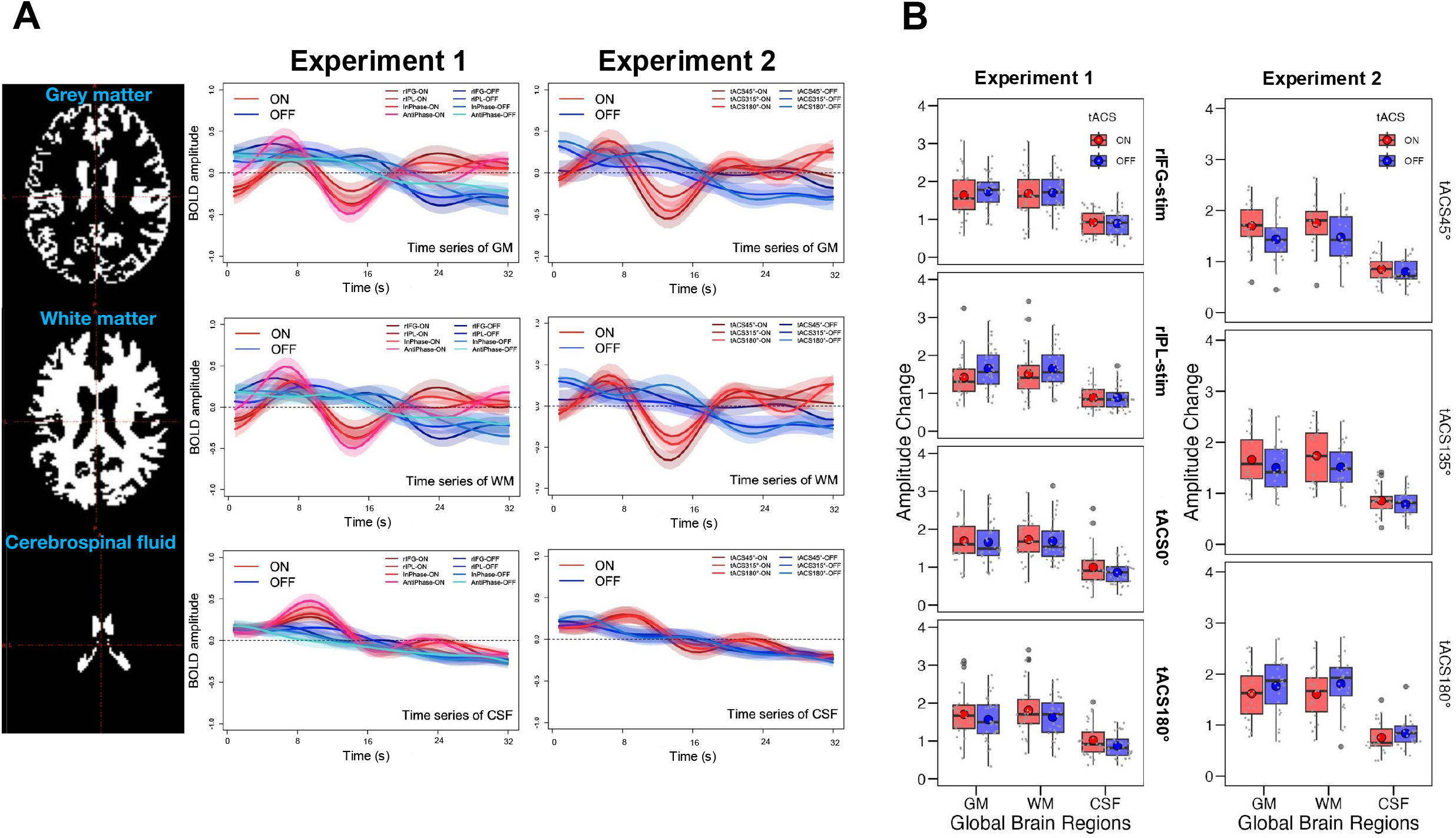
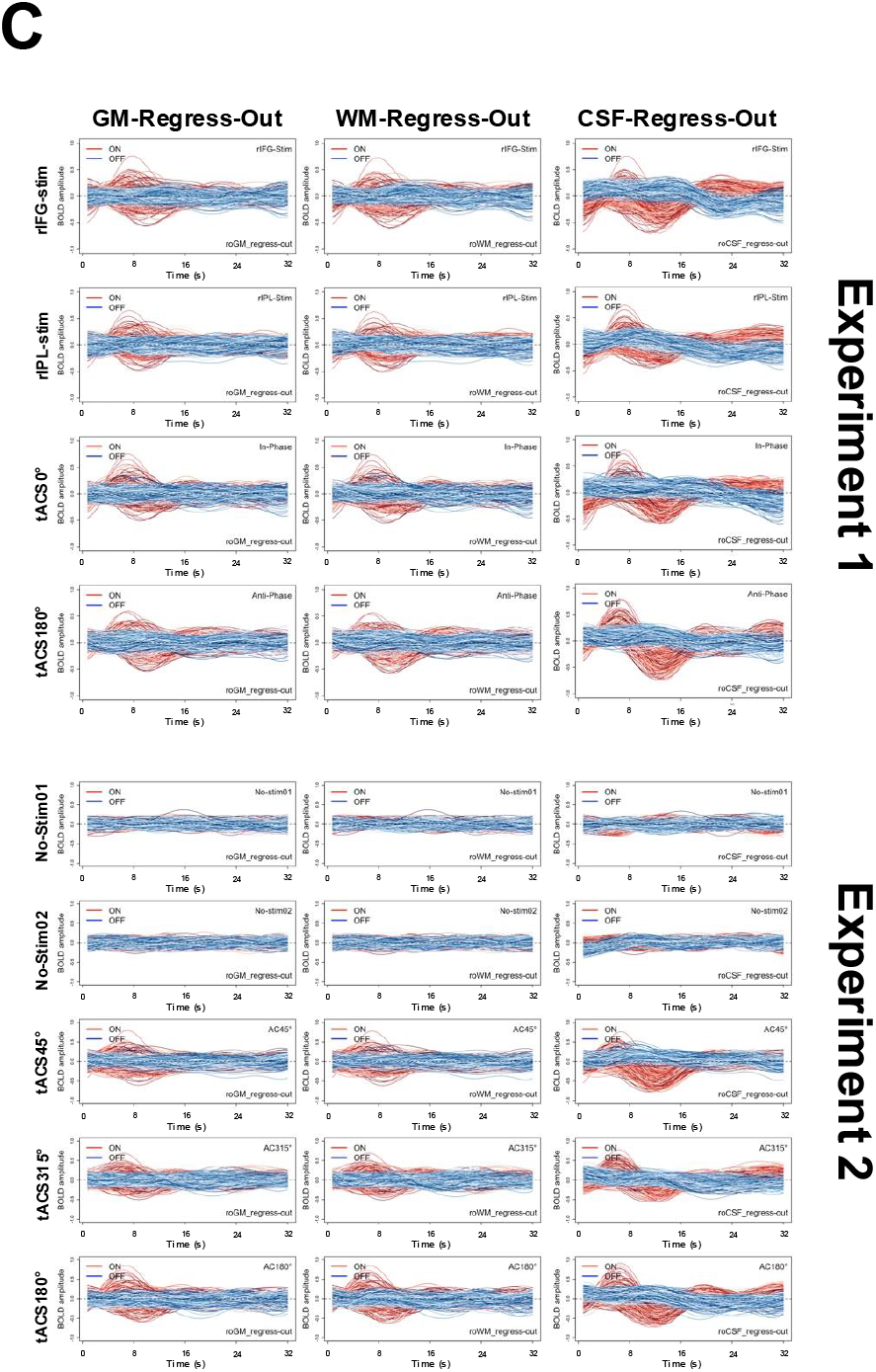
The effects of tACS on global brain regions: GM, WM, and CSF. (A) Each graph illustrates the voxel-wise averages of the BOLD signals within the GM, WM, and CSF for each tACS condition in Experiments 1 and 2. The averages are calculated separately for tACS-ON and -OFF epochs. The ribbons in each graph represent mean ± SE. (B) The box plots illustrate the mean change in BOLD signal during the tACS-ON and -OFF epochs for each tACS condition across experiments. (C) Depicts the results of averaging across tACS-ON and -OFF epochs for each ROI defined by Power ROI. The BOLD signals within each ROI are shown after regressing out signals from the GM, WM, and CSF. The upper panel corresponds to Experiment 1, while the lower panel corresponds to Experiment 2. tACS, transcranial alternating current stimulation; GM, grey matter; WM, white matter; CSF, cerebrospinal fluid; BOLD, blood oxygenation level-dependent; ROI, region of interest.

Two-way repeated measures ANOVA (tACS conditions x ON/OFF x anatomical subdivision) showed the significant main effect of anatomical subdivision (Experiment 1: *F*(2,62)=222.36, *p*<.001, *η*_*p*_^*2*^=.88; Experiment 2: *F*(2,46)=260.57, *p*<.001, *η*_*p*_^*2*^=.92) and the marginally significant interaction between tACS conditions and ON/OFF only for Experiment 2 (*F*(2,46)=1.24, *p*=.06, *η*_*p*_^*2*^=.12). Post-hoc comparison following the main effect of anatomical subdivision revealed that, in Experiment 1, amplitude change in the CSF was lower than that in the GM (*t*(31)=14.54, *p*<.001) and WM (*t*(31)=15.63), *p*<.001), and amplitude change in the WM was greater than that in the GM (*t*(31)=3.82, *p*=.0006). In Experiment 2, amplitude change in the CSF was lower than that in the GM (*t*(23)=16.38, *p*<.001) and WM (*t*(23)=17.10, *p*<.001), and there was a marginally significant difference between amplitude changes in the GM and WM, showing that amplitude change in the WM was greater than that in the GM (*t*(23)=1.81, *p*=.08). Regarding the interaction between tACS conditions and ON/OFF, a simple main effect of tACS conditions at OFF was significant (*F*(2,46)=3.77, *p*=.03, *η*_*p*_^*2*^=.14), and there were marginally significant differences across tACS conditions, showing that the amplitude change in tACS 180° at the tACS-OFF epoch was higher than that in tACS 45° (*t*(23)=2.46, *p*=.06) and tACS 135° (*t*(23)=1.99, *p*=.06), whereas there was no significant difference between tACS 45° and tACS 135° (*t*(23)=0.38, *p*=.71).

Additionally, we regressed the GM, WM, and CSF from the time series of each ROI in the whole brain and compared the tACS-ON and -OFF epochs (Figure 6C). Consistent with the results above, the most prominent effect of tACS-induced signal elevation was observed in the time series, with the CSF regressing out consistently across the experiments. In other words, the effects of tACS, which were previously considered to only have localised effects in the cerebral cortex, are likely to propagate across broad regions of the brain through the fasciculus.

### Propagation of BOLD signals across whole brain

As demonstrated in the Supplementary animation, the stimulation effects seemed to propagate from the stimulated region and the corresponding contralateral area to encompass the whole brain. Hence, the question of qualifying how stimulation effects propagate across the brain arises. To examine the propagation of BOLD signals across the whole brain, the 264 ROIs were sorted based on the peak values of BOLD signals, and the results were visualised in heatmaps (Figure 7).

**Figure 7.**
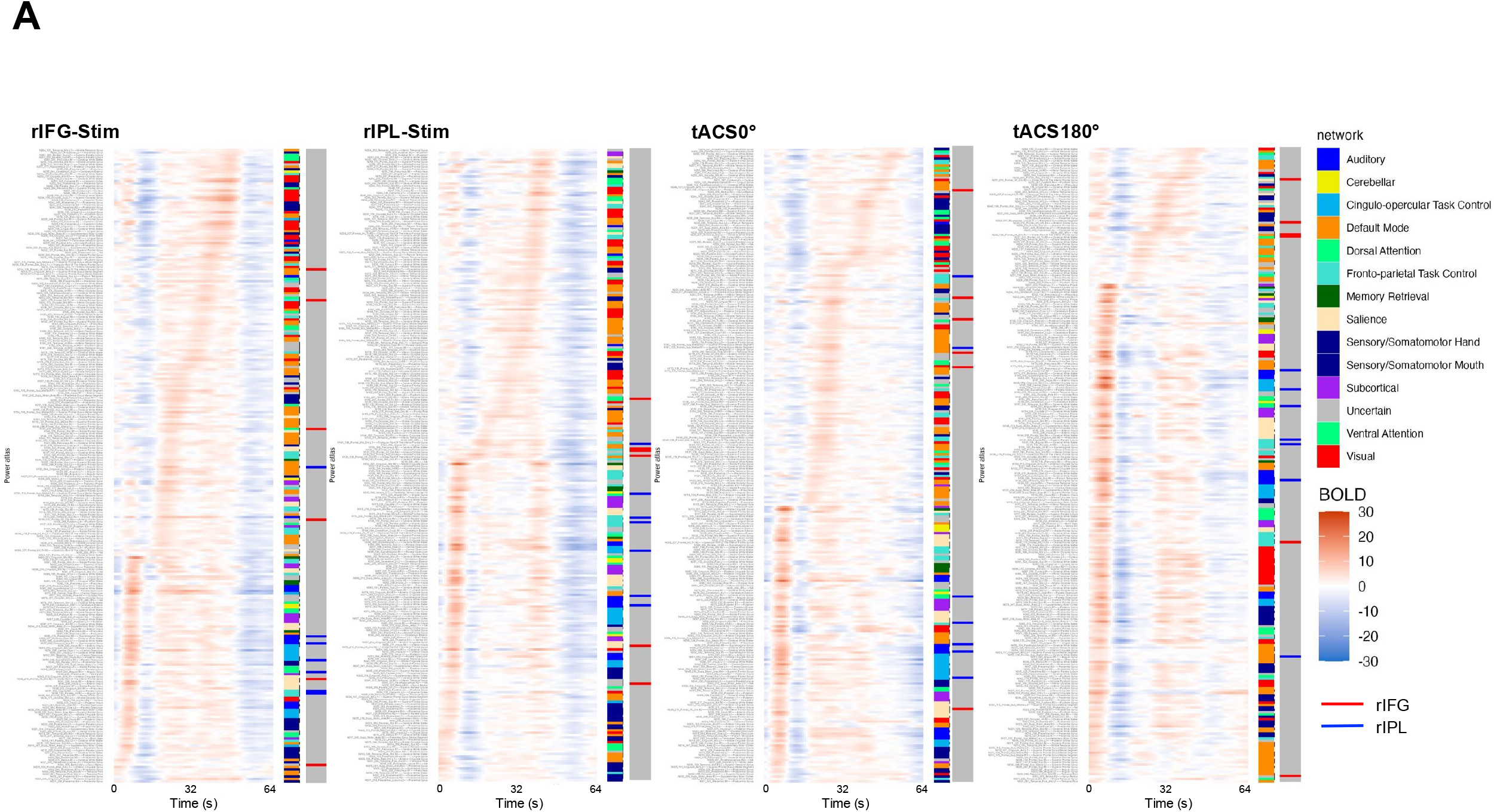
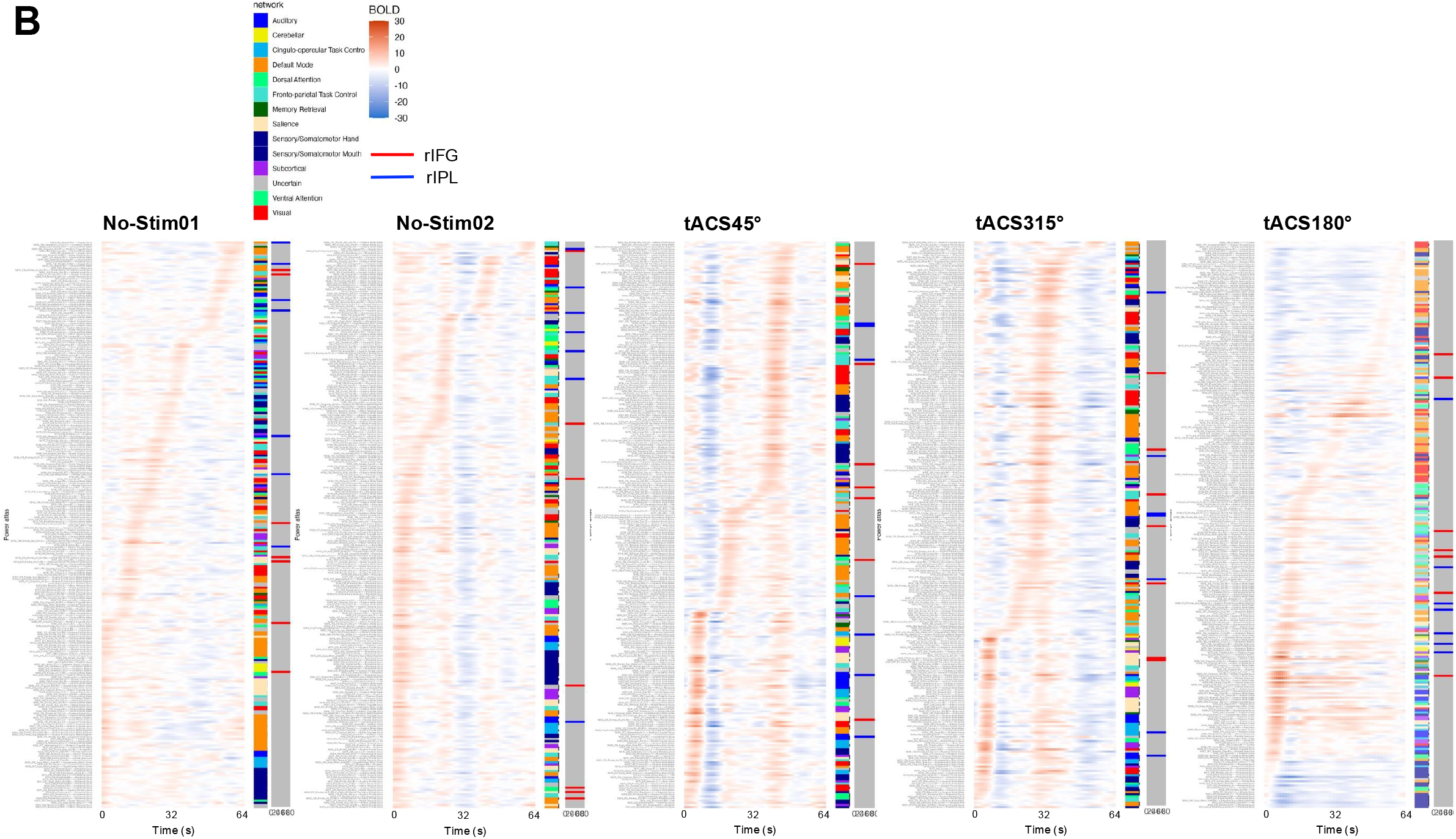
Propagation of the BOLD signals across the whole brain. (A) and (B) illustrate the heatmaps containing the BOLD signals of each ROI in the whole brain for Experiments 1 and 2. The signals were averaged across consecutive tACS-ON and -OFF epochs and sorted according to the peak timing of each ROI. The two-coloured bars on the right side of each heatmap represent the functional networks defined by the Power ROI on the left and the stimulated regions on the right. BOLD, blood oxygenation level-dependent; ROI, region of interest.

When comparing the no-stimulation and stimulation conditions, it was evident that the range of signal value fluctuations were small in the no-stimulation condition (No-Stim01 and No-Stim02 in Figure 7B). Under no-stimulation conditions, the sorting of ROIs by BOLD signal peak values generated almost random arrangements of the ROIs according to the network organisation (colours randomly changed in the network column). In contrast, the tACS conditions exhibited a relatively consistent pattern of network organisation in Experiments 1 and 2. We then elaborated on the network organisation based on the sorting of ROIs by the peak values of the BOLD signal. Under the tACS conditions, the somato-motor network firstly exhibited peak values. Subsequently, a transition of peaks was observed in the order of the frontoparietal, dorsal/ventral attention, visual, and default mode networks. Although there were instances where a clear organisation of networks was not evident under stimulation conditions, a consistent pattern was evident compared to the no-stimulation conditions.

Qualitatively, under no-stimulation conditions, variability in the timing of the peak BOLD signal values for both the rIFG and rIPL was observed (Figure 7B). In contrast, during Experiment 1, the rIPL regions first exhibited peaks, followed by the rIFG regions. In Experiment 2, under the tACS 45° and tACS 315° conditions, there was no clear trend in the peak timing of the regions near the rIFG and rIPL. However, under the tACS 180° condition, a tendency in which the rIPL region exhibited initial peaks followed by the rIFG region was observed.

## Discussion

The objectives of the present study were three-fold: 1) to elucidate the immediate online effect of tACS by comparing the short-term BOLD signals during tACS with periods where no stimulation is applied, 2) to examine the extent of the influence on the brain when applying tACS, and 3) to explore whether variations in the phase difference between two brain regions result in differential effects on the stimulated areas and the whole brain. Based on these objectives, we identified three key findings. First, immediate online alterations in the BOLD signal of the stimulated region during the periods where tACS was applied. Second, the online tACS effect resulted in pronounced changes in BOLD signals, particularly in regions rich in neurons, such as the GM and WM, while the BOLD changes in CSF were minimal. Third, stimulation of the two regions with a specific phase difference resulted in a tendency of an increase in the BOLD signal, with a temporal lag in the peak signal values of the stimulated region.

The immediate online changes observed in the resting-state BOLD signals due to tACS in this study contribute to a refined understanding of the previously documented BOLD signal modulation effects observed during task-related activities (Chai et al., 2018; Violante et al., 2017; Vosskuhl et al., 2016). The BOLD signal modulation effects of tACS during rest have not been clearly demonstrated, and it was previously considered that these effects were only evident during tasks in which specific brain functions were activated (Vosskuhl et al., 2016). However, our findings demonstrate that tACS applied to single and dual regions induces online effects, even during the resting state. Statistical analysis revealed that the behaviour of BOLD signals during tACS-ON epochs differed depending on the stimulated regions when compared to epochs without stimulation (tACS-OFF). In the rIFG region, significant increases were observed during the latter half of the tACS-ON epochs for both the rIPL single-site stimulation and the other dual-site stimulation conditions compared with the signals in tACS-OFF epochs. However, no significant increase was observed in the rIFG single-stimulation condition. Conversely, in the rIPL region, there was a waveform change indicating an increase in BOLD signals during the first half of the tACS-ON epochs, but statistically significant differences compared to tACS-OFF epoch BOLD signals were observed only in the tACS 180° condition in Experiment 1. These results suggest that although the effects of tACS were minimal during tACS-OFF epochs, they might still persist to some extent, potentially indicating incomplete return to a normal state of the BOLD signal during resting.

Consequently, to examine changes in BOLD signals, a correction was applied using tACS-OFF epoch as a baseline immediately before the tACS-ON epochs. From the results of the BOLD signal during the tACS-ON epochs in the target regions (Figure 4), it can be observed that in conditions where the rIFG was stimulated, the BOLD signal of the rIFG started to decline with the onset of tACS. Upon comparison with the baseline, significant increases were observed in the rIFG region during the latter half of the tACS-ON epochs in Experiment 1 under the tACS 180° condition, as well as significant decreases during the first half of the tACS-ON epochs in Experiment 2 under the same condition. Additionally, significant decreases were observed under the tACS45° condition. Conversely, in the rIPL region, significant increases in BOLD signals during the first half of the tACS-ON epochs were observed only under the tACS 180° condition in Experiment 2. These results suggest a potential asymmetry, particularly indicating an immediate decrease in BOLD signals in the rIFG and an increase in the rIPL under the tACS 180° condition. Qualitatively, without stimulation to the rIPL, the BOLD signal of the rIFG did not decline. In the case of rIFG single-site stimulation, the rIFG signal did not fall below the baseline of zero, whereas in the two-region stimulation condition, it dropped below zero. This may suggest that, even when stimulating the same region, there are differences in the effects of tACS between single- and dual-site stimulations. In contrast, the rIPL signal exhibited an immediate increase in all conditions, including the rIFG stimulation condition, where no stimulation was applied to the rIPL region, which differed from the results of the rIFG signal. From the tACS results of a single region, it is more likely that rIFG and rIPL are regions with mutual interactions rather than regions with solely independent functioning. These results align with reports on the effects of tACS based on intracranial neural activity measurements in non-human primates (Berényi et al., 2012; Johnson et al., 2020; Krause et al., 2019, 2022), supporting the broader applicability of tACS beyond task-related activation. Given such an interaction, the subsequent interpretation of the results will focus on elucidating the extent to which the tACS effects occur within the whole brain.

Regarding the extent of the stimulation effects within the whole brain, the observed modulation of BOLD signals within the stimulated region aligns with findings from previous studies. Several studies have demonstrated that tACS with frequencies specific to the visual cortex elicits an increase in BOLD signals compared with sham stimulation (Chai et al., 2018; Vosskuhl et al., 2016). Previous investigations have primarily focused on changes within the stimulated region, neglecting the assessment of alterations in other brain regions. In this study, we examined BOLD signals beyond the stimulated area. While BOLD signal modulation in the stimulated region was pronounced compared to that in other areas, a similar trend of modulation was evident in non-stimulated regions with different peak timings from those in the stimulated regions (Figure 5). Upon investigation of the online effects of tACS by dividing the whole brain into the GM, WM, and CSF, notable online effects of tACS were observed specifically in the GM and WM compared with the CSF (Figure 6A). In both Experiments 1 and 2, the amplitude change of BOLD in tACS-ON and -OFF epochs was shown to be greater in the GM and WM compared with that in the CSF. Furthermore, it was found that the amplitude change in the WM was larger than that in the GM. Specifically, in the tACS-OFF epochs of Experiment 2, the amplitude change of BOLD was greater in the tACS 180° condition compared to the 45° and 135° conditions. These results suggest that the effects of localised stimulation extend to a broader scope within the whole brain through neural fibre connections (Thiebaut de Schotten et al., 2011). Given that the rIFG and rIPL targeted with tACS in this study anatomically communicate via WM tracts (i.e., the superior longitudinal fasciculus), the effects of tACS may be particularly pronounced in the WM. It is plausible that the stimulatory effects are propagated throughout the brain from these two regions via this structural linkage.

The influence of tACS extends across broad regions of the brain, including the GM and WM. However, the question of how the impact of tACS propagates throughout the brain remains. Sorting the 264 regions of the whole brain based on the peak values of the BOLD signal revealed that the effects of tACS were spread across functional brain networks (Figure 7). In contrast, under the no-stimulation condition, there was no cohesive organisation at the network level, and the spread was random. An essential finding was the transition of peak values in stimulation conditions, starting with the sensorimotor network and subsequently transitioning through the frontoparietal region, including the stimulated region, attention, visual, and default mode networks. Although there were slight variations in the order across experiments, the consistent observation of transitions in peak values under stimulation conditions provides insights into how tACS affects the cascade through different functional brain networks. While previous studies have hinted at the network-level effects of tACS (Cabral-Calderin et al., 2016; Preisig et al., 2021; Weinrich et al., 2017), they were limited to specific networks, and systematic transitions across the entire brain were not explored.

The results suggest that the sensorimotor regions are initially activated, likely arising from sensory evocation due to stimulation. Subsequently, the activity spread from the stimulated adjacent regions, involving anterior-posterior and lateral-medial transitions, were noted. It is conceivable that, by inputting stimuli that activate specific networks, transitions occur in the overall brain state (peak value transitions along with functional networks). Further examination of the results regarding the order of peak activation timing in the stimulated regions, as illustrated in Figure 7, showed that in the 180° phase-difference condition of Experiments 1 and 2, the nearby regions of the rIPL first reached their peak, followed by peaks in neighbouring regions of the rIFG. This finding aligns with the temporal changes in the time series of the stimulated regions shown in Figure 3, suggesting an effect of phase-differentiated tACS, especially at a phase difference of 180°, in transitioning peak brain activity from the parietal to frontal regions. The transition of activity peaks along the network and temporal changes in local regions within this transition, induced by phase-differentiated tACS, are likely rooted in the spatiotemporal dynamics of both local and global neural activity. Moreover, simulation study has not only suggested that the distribution of the electric field in the brain depends on the phase difference (Saturnino et al. 2017) but also that the electric field in the anti-phase is about twice that in the in-phase. Therefore, the specific transition pattern observed in the present study under the 180° phase difference condition is consistent with these previous studies (Alekseichuk et al. 2019).

While this interpretation needs to be quantitatively validated, it raises the possibility that tACS not only induces modulation of activity in localised regions but also leads to changes of specific functional patterns (i.e., functional networks, brain states, and neural manifolds). In recent years, an approach has been reported that shifts the focus from local brain activity to global spatiotemporal dynamics, capturing the essence of the brain by reframing overall brain activity as transitions in the state space (Asai et al., 2023). In this sense, future tES research should aim to clarify both the local effects in stimulated regions and the subsequent effects that propagate to other regions. These clarifications are consistent with findings supporting functional localisation in stimulated regions, while also suggesting the need for a more integrative perspective when examining broader brain dynamics (Barack & Krakauer, 2021).

## Conclusions

We demonstrated an online modulation effect on the BOLD signal through tACS, revealing that this effect, specifically evident in the dual-site anti-phase tACS condition, extends not only to the locally stimulated area but also propagates across the whole brain in accordance with anatomical features. Furthermore, the propagation of the stimulation effect throughout the whole brain followed functional networks, and concurrently, within the stimulated region, a transition was observed from the parietal to the frontal regions. Thus, tACS with a specific phase difference in two anatomically connected brain regions can immediately modulate online neural dynamics at both local and global scales.

## Limitations and correction of the current study

This study has several limitations. First, the temporal synchronisation between tES output and fMRI acquisition relied on manual triggering after a nominal ten dummy-scan delay (8 s with a TR of 0.8 s). However, post-hoc verification revealed that scanner latency made the dummy period slightly different from 8 s, and manual initiation introduced a human reaction delay. Collectively, these factors potentially caused a sub-second offset in stimulation onset and a minor temporal shift when averaging the ON/OFF cycles. Future studies should use an automatic stimulation control system to enable precise synchronisation between the MRI trigger signal and tES output, for example, via MATLAB-based trigger detection and device control, thereby ensuring temporal accuracy.

Second, the stimulation intensity reported initially in this study, a peak-to-peak amplitude of 1 mA, should be corrected to 4 mA (see Methods). This discrepancy arose from an inconsistency between the impedance-monitoring device manual and the stimulator’s actual hardware design. Although the delivered current intensity remained within established safety limits (Antal et al., 2017; Bikson et al., 2016) and no adverse events were observed, the higher current would have increased current density at the electrode– skin interface and cutaneous sensation relative to the originally reported 1 mA, suggesting that somatosensory cues were potentially more salient than initially described.

Third, interpretations of the BOLD effects should consider that the observed temporal modulation, both near the stimulated regions and across the whole brain, may arise partly from activation of cutaneous afferents or peripheral nerves, rather than solely from cortical entrainment. Previous studies have reported tES-induced activation of the insula and somatosensory regions, including Cabral-Calderin et al. (2016); however, the relationship between these activations and perceived stimulation sensations remains unclear. Given the substantial current attenuation by the scalp and skull (Vöröslakos et al., 2018) and the ongoing debate about whether tES effects are mediated by direct cortical modulation or by peripheral sensory input (Asamoah et al., 2019), the present findings cannot be interpreted as purely reflecting brain stimulation effects. Therefore, the possibility that cutaneous sensations contributed to the results cannot be excluded, and all effects should be interpreted with appropriate caution.

## Acknowledgements

This work was supported by JSPS KAKENHI (Grant Numbers 20J00669 to KH, 19H05725 to HI, 19H01777 to TA and HI, 21H00959 to HI, TA, and KH) and the Innovative Science and Technology Initiative for Security (Grant Number JPJ004596, ATLA, Japan to TA and KH). We thank Chimoto Takeuchi, Ayako Tsukamoto, Masaru Tanaka, and Takamasa Hamamoto for their support in the experiments. We appreciate the support of the WPI-IRCN Human fMRI Core at the University of Tokyo Institute for Advanced Studies.

